# Arabidopsis DXO1 is an evolutionarily diverged homolog that impacts ribosome loading and mRNA surveillance

**DOI:** 10.1101/2025.10.29.684918

**Authors:** Diep R. Ganguly, Xiang Yu, Matthew D. Mortimer, Marten Moore, Pravin Khambalkar, Barry J. Pogson, Brian D. Gregory

**Affiliations:** Department of Biology, University of Pennsylvania, Philadelphia, PA, 19104, USA; School of Life Sciences and Biotechnology, Shanghai Jiao Tong University, Shanghai, China; Research School of Biology, The Australian National University, Canberra, ACT, 2601, Australia

**Keywords:** DXO1, mRNA surveillance, evolutionary analysis, degradome, ribosome loading, translation, Arabidopsis thaliana

## Abstract

The Rai1/Dxo1/DXO protein family are involved in eukaryotic 5′-end RNA quality control by hydrolyzing non-canonical mRNA cap structures, such as the NAD^+^ 5′-cap. The plant DXO1 ortholog shows distinct biochemical properties, compared to its fungal and mammalian counterparts, and only minor phenotypic traits can be attributed to its NAD^+^-decapping (deNADding) activity. On the other hand, the chloroplast and growth defects observed in *dxo1* null mutants appear linked to the presence of a serine-rich domain at its N-terminus. Here, we report that plant DXO1 is a distinct ortholog that likely diverged earlier than its fungal and mammalian counterparts. We attribute this to the presence of a serine-rich N-terminal domain, which is found almost exclusively in plants and likely arose during the emergence of angiosperms. Degradome profiling revealed that Arabidopsis DXO1 contributes to mRNA surveillance, via examination of exon junction complex and ribosome footprints, and this activity is reliant on the presence of the serine-rich N-terminal domain in Arabidopsis. Furthermore, polysome profiling revealed that plants lacking DXO1 have severe ribosome loading defects, which may explain the loss of mRNA surveillance and alterations in pre-rRNA processing. Taken together, our data leads us to propose a model whereby Arabidopsis DXO1 is involved in 5′-end RNA quality control, which is critical for ribosome loading that impacts translational fidelity and translation-dependent mRNA surveillance.

## Introduction

The Rai1/Dxo1/DXO family of enzymes are implicated in 5′-end RNA quality control, due to their ascribed core NAD^+^-cap hydrolase (deNADing) activity alongside ortholog-specific functions as a (pyro)phosphohydrolase, hydroxyl dinucleotide hydrolase, FAD cap hydrolase (deFADding), and/or 5′-to-3′ exoribonuclease (1–6). Work in fungi highlighted that the multifaceted activity of yeast Rai1 allows for the hydrolysis of incomplete and non-canonical mRNA cap structures, releasing a 5′-monophosphorylated RNA that can be degraded by the major nuclear 5′-exoribonuclease, Rat1, with which it interacts (6, 7). Genome duplication within the Saccharomyces lineage produced a Rai1 paralog, Dxo1, which acts as a distributive exoribonuclease that enables 5′-end processing of incompletely capped RNA and 25S rRNAs by the cytoplasmic 5′-exoribonuclease (XRN), Xrn1 (4, 8). Mammalian DXO is structurally similar to both yeast proteins, Rai1 and Dxo1, and harbors shared properties such as pyrophosphohydrolase, decapping, exoribonuclease, deNADding, and deFADding activity, but does not interact with the mammalian Rat1 homolog, XRN2 (3, 6, 7). Arabidopsis harbors a single copy of DXO1 that is most homologous to mammalian DXO. Arabidopsis DXO1 lacks the domain required for interaction with plant XRNs and contains amino acid substitutions that limits its activity to deNADding (1, 2, 9). It also harbors an intrinsically disordered and serine-rich N-terminal domain (NTD), which is considered to be plant-specific (1, 2). Therefore, while the Rai1/Dxo1/DXO protein family is conserved among eukaryotes, there appear to be distinct functions between fungal, mammalian, and plant homologs. The evolutionary relationship between Rai1/Dxo1/DXO homologs is yet to be systematically evaluated, especially in the context of the prevalence of the serine-rich NTD.

Despite extensive biochemical characterization of Rai1/Dxo1/DXO proteins *in vitro*, there is an incomplete understanding of their *in vivo* impacts. Work in Arabidopsis suggests that DXO1 degrades NAD^+^-capped RNAs, many of which encode ribosomal proteins, which are otherwise processed into small RNAs (9). This is consistent with DXO1 functioning in 5′-end RNA quality control for transcriptome integrity. Furthermore, in plants, NAD^+^-capping has been positively associated with both mRNA degradation (9) and ribosome association (10), highlighting how deNADing could impact mRNA regulation. However, the severe developmental defects and malformed chloroplasts observed in plants lacking DXO1 have been associated with a deNADding-independent function (1, 2). Instead, deletion of its serine-rich NTD recapitulates the developmental defects in *dxo1* null plants, alongside the mis-expression of ribosomal genes and accumulation of rRNA precursors (pre-rRNAs) (11). This serine-rich NTD facilitates an interaction with RNA GUANINE-7 METHYLTRANSFERASE 1 (RNMT1) in the nucleus, leading to its activation to catalyze the formation of the 7-methylguanosine (m^7^G) mRNA cap (12). In this way, DXO1 and RNMT1 appear to work in tandem to provide a plant-specific counterpart of RNMT and RNMT-Activating Mini protein (RAM), which jointly perform cap methylation in mammals (13). Collectively, these observations broaden the potential impacts of DXO1 on mRNA regulation, independent of its deNADding activity, which are yet to be identified.

Two important forms of post-transcriptional regulation include mRNA degradation and ribosome loading (14–16). In plants, the global profiling of mRNA degradation intermediates (the ‘degradome’) by capturing RNAs harboring 5′-monophosphate (5′-P) ends, representing fragments undergoing XRN-mediated 5′-to-3′ decay, has facilitated novel insights on the regulation of multiple mRNA decay pathways (17). For example, the detection of 5′-P ends mapping to regions predicted to be occupied by exon-junction complexes (EJCs, EJC footprints) and terminating ribosomes (ribosome footprints). These footprints reflect RNA decay that occurs before, or during, the pioneer round of translation (18) or co-translationally (co-translational RNA decay, CTRD) (19, 20), respectively. Since both processes follow at least one round of translation, these signals can be collectively considered to reflect translation-dependent mRNA surveillance-triggered decay (14, 16).

Converse to mRNA decay is ribosome loading–the recognition of 5′-m^7^G capped mRNAs by eukaryotic initiation factor (eIF) 4E leading to the assembly of the eIF4F complex, which recruits the pre-initiation complex that scans for the start codon after which the complete 80S ribosome is formed (21). Changes in ribosome loading entails the formation, or dissociation, of mRNA-ribosome complexes (polysomes), which can impact both the ribosome occupancy (fraction of mRNAs with at least one ribosome) and ribosome density (number of ribosomes on an mRNA) to influence the translation rate for a given pool of transcripts (22, 23). Global changes in ribosome loading have been routinely diagnosed via polysome profiling, which has revealed the dynamic nature of polysome formation and dissociation in response to developmental, circadian, and environmental factors (15, 24).

Here, we provide evidence that Arabidopsis DXO1 is an evolutionarily diverged homolog that impacts mRNA surveillance-triggered decay, ribosome loading, and rRNA processing. First, we highlight that angiosperms harbor a divergent Rai1/Dxo1/DXO homolog that is characterized by a serine-rich NTD, which is largely absent in non-flowering plants, metazoa, and fungi. Degradome analysis revealed that lack of Arabidopsis DXO1 results in elevated EJC footprints and loss of CTRD footprints, indicative of impaired mRNA surveillance-triggered decay. Analysis of transgenic lines expressing DXO1 variants (1) links this to the presence of the serine-rich NTD, positing a specialized function of this homolog in angiosperms. Integrating our degradome data with rRNA annotations and polysome profiling revealed that loss of DXO1 affects pre-rRNA processing and severely impairs ribosome loading. Taken together, our findings lead us to propose that 5′-end RNA quality control by Arabidopsis DXO1 promotes ribosome loading, which is required for translation-dependent mRNA surveillance-triggered decay.

## Results

### Angiosperm DXO1 shows evolutionary divergence from non-flowering plant, fungal, and animal homologs

We performed an evolutionary analysis to explore the ancestry between Rai1/Dxo1/DXO homologs, hereafter referred to as DXO1 homologs. A sequence similarity network of Pfam clan CLO236 yielded no detectable prokaryotic orthologs (E-value ≤ 1), suggesting that this family has diverged extensively within eukaryotes. We then retrieved sequences from the Pfam family (PF08652) containing Arabidopsis DXO1 (AT4G17620.1), which underwent manual curation and filtering to generate a high-confidence, non-redundant dataset (**Supplementary Table 1, Supplementary Figure 1A**). With this, we generated a cross-kingdom K-Nearest Neighbors (KNN) map (**Figure 1A**). This showed that most plant DXO1 sequences are distinct from animal and fungal counterparts, although some sub-groups cluster within metazoa and fungi. This suggests that most plant sequences are evolutionarily distinct while some resemble fungal and metazoan homologs. Therefore, plant-like DXO1 likely diverged after the separation of Archaeplastida from Opisthokonts. We investigated this further using the evolutionary velocity pseudo-time tool, which facilitates the prediction of the root sequence(s) (25). Two likely roots were identified, one within fungi, which appears to be more similar to metazoan sequences, and another within Viridiplantae (**Figure 1B**). This makes sense given the absence of a common prokaryotic ancestor, while also aligning with biochemical evidence that Arabidopsis DXO1 is distinct from fungal and metazoan homologs.

**Figure 1.**
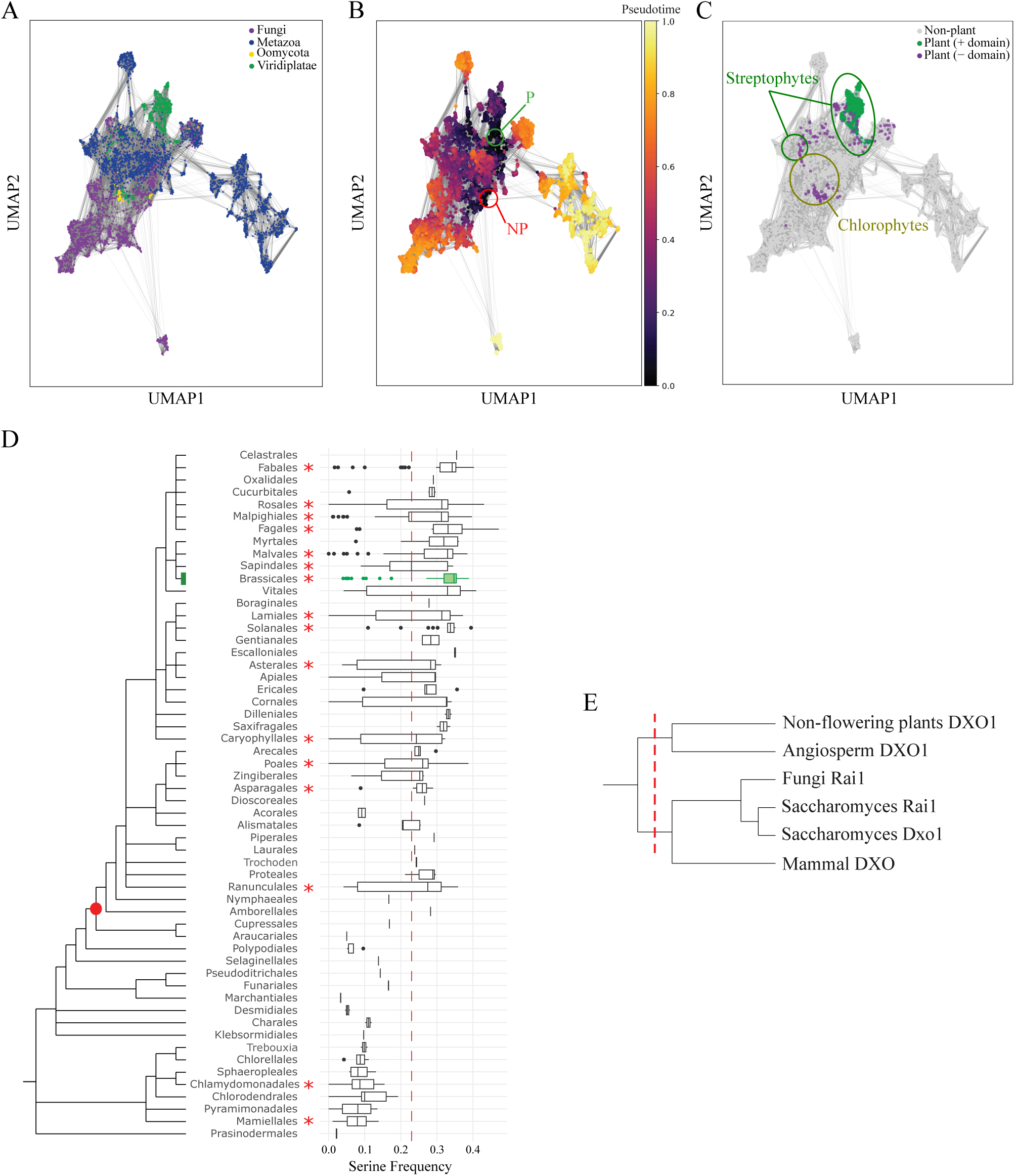
Angiosperms harbor an evolutionarily divergent DXO ortholog characterized by a serine-rich N-terminal region. (A) K-Nearest Neighbours (KNN) network of DXO orthologs generated using Uniform Manifold Approximation and Projection (UMAP) with 25 nearest neighbours. Points are colored by taxonomic kingdom. (B) Evolutionary velocity pseudo-time plot, with sequences colored along a gradient from 0 (ancestral-like) to 1 (most diverged from ancestral-like). Two ancestral-like regions are marked as the predicted roots for plant (P) and non-plant (NP) orthologs. (C) KNN plot from (A) recolored to highlight plant sequences with (green) and without (purple) a serine-rich N-terminal domain. Non-plant sequences are shown in grey. Streptophyte and chlorophyte sequences are labeled; unlabeled plant sequences represent mixed clades. (D) Cladogram of plant orders with accompanying box-and-whisker plots depicting serine frequency. The Brassicales branch is highlighted in green. Red node indicates the divergence of angiosperms. Red dotted line marks the threshold used to define the presence of a serine-rich N-terminus. Serine frequency was calculated from the first 18% of the N-terminal region of each sequence. Red stars denote clades with sample sizes ≥ 10. Sample counts for each clade are provided in Supplementary Table 1. (E) Order-level cladogram of DXO1 and DXO1-like homologs. The vertical dashed red line indicates the cut-off from the last common (prokaryotic) ancestor, defining two pseudo-roots within the dataset.

One of the distinguishing features of Arabidopsis DXO1 is its serine-rich NTD. We evaluated the presence or absence of this domain by calculating serine frequency within the first 18% of the amino acids for each homolog. This threshold was based on manual curation of an alignment identifying this domain using Arabidopsis as a reference. We observed a bimodal distribution in serine residue frequency where a value of 0.23 was determined as the trough, arising from a subset of Viridiplantae sequences (**Supplementary Figures 1B-C**). Therefore, we considered DXO1 homologs with serine frequency > 0.23, within the first 18% of amino acids, as harboring an Arabidopsis-like serine-rich NTD (**Supplementary Table 1**). We detected this domain almost exclusively in plants (**Figure 1C**), with the exception of eight non-plants sequences (**Table 1**, **Supplementary Figures 1D**). Further investigation revealed that this pan-grouping comprises Streptophyta, with a small group of streptophytes clustering with metazoa sequences. In fact, the serine-rich NTD appeared to be limited to a subset of Streptophyta, suggesting that it arose within this clade. To examine this further, we constructed a cladogram co-plotted with the N-terminal serine frequency for sequences grouped by plant order (**Figure 1D**). This suggests that the serine-rich NTD emerged alongside the divergence of Angiosperms. Notable is the presence of Angiosperm clades with sequences without a serine-rich N-terminus. This could be explained by gene annotation errors, or multiple gain- or loss-of-function events. Given its prevalence across all streptophyte orders, and that the Streptophyta sequences without the serine-rich NTD predominately cluster with the metazoan sequences, we suggest a null-hypothesis of loss-of-function.

**Table 1.** Non-plant sequences with serine-rich N-terminal domain.

| UniProtKB ID | Kingdom | Phylum | Class | Order | Serine Frequency |
| --- | --- | --- | --- | --- | --- |
| A0A316ZA69 | Fungi | Basidiomycota | Exobasidiomycetes | Entylomatales | 0.24 |
| A0AAN6GCK9 | Fungi | Basidiomycota | Exobasidiomycetes | Tilletiales | 0.29 |
| A0AAD5TBT3 | Fungi | Chytridiomycota | Chytridiomycetes | Spizellomycetales | 0.27 |
| A0A8S1F7G5 | Metazoa | Nematoda | Chromadorea | Rhabditida | 0.24 |
| A0AA36BQR8 | Metazoa | Mollusca | Cephalopoda | Octopoda | 0.36 |
| A0A267DQM7 | Metazoa | Platyhelminthes | Rhabditophora | Macrostomida | 0.35 |
| A0A267EM17 | Metazoa | Platyhelminthes | Rhabditophora | Macrostomida | 0.36 |
| A0A267F4U8 | Metazoa | Platyhelminthes | Rhabditophora | Macrostomida | 0.36 |

### DXO1 promotes translation-dependent mRNA surveillance-triggered decay independently of its deNADding activity

Based on our previous analyses (9), we hypothesized that DXO1 could impact mRNA surveillance-triggered decay. To explore this, we utilized global mapping of uncapped and cleaved transcripts (GMUCT) data on 12-day-old WT and *dxo1* seedlings (GSE142388, **Supplementary Table 2**) to compare EJC and ribosome (or CTRD) footprints (18, 19). This re-analysis revealed that *dxo1* seedlings had elevated EJC footprints and suppressed ribosome footprints, suggestive of increased decay prior to the pioneer round of translation and limited CTRD, respectively (**Figures 2A-B**). We tested whether this was a consequence of increased ribosome stalling by incorporating GMUCT data (GSE71913) from leaves treated with cycloheximide (CHX) (19), which induces ribosome stalling by blocking the E-site (26, 27). CHX treatment resulted in a small reduction in EJC footprints and almost completely abrogated CTRD footprints, with pronounced 3-nt periodicity upstream of the stop codon. Importantly, these footprints are distinct from those observed in *dxo1* seedlings, especially in the case of EJC footprints. Together, this suggests that DXO1 function impacts mRNA surveillance without causing ribosome stalling.

**Figure 2.**
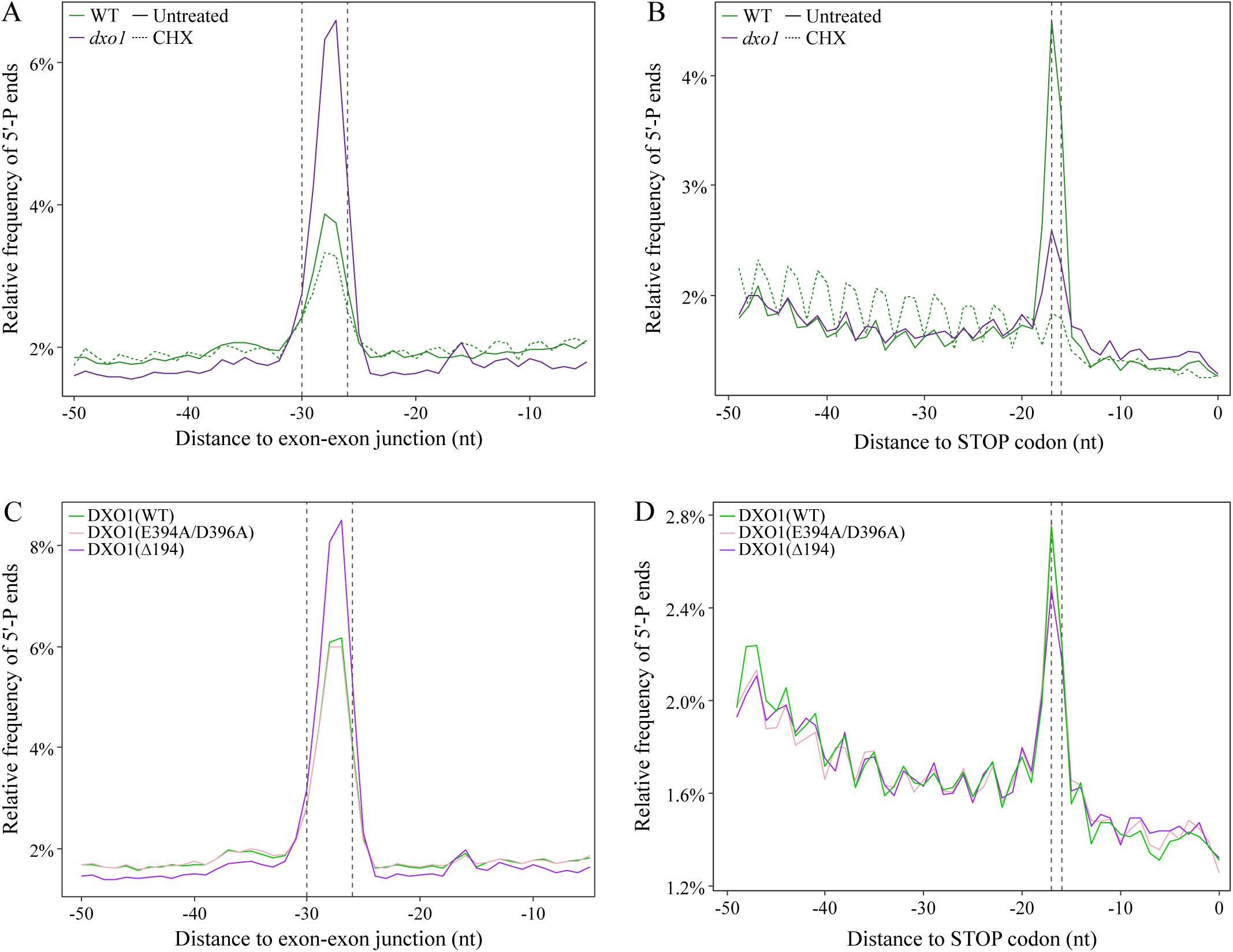
DXO1 impacts EJC footprints and co-translational RNA decay through its N-terminus. (A-B) Relative frequency of 5’-P ends upstream of (A) exon-exon junctions and (B) stop codons in WT, untreated and cycloheximide (CHX)-treated, and *dxo1* plants. (C-D) Relative frequency of 5’-P ends upstream of (C) exon-exon junctions and (D) stop codons in DXO1(WT), DXO1(E394A/D396A), and DXO1(Δ194) transgenic Arabidopsis lines. Dotted lines represent the expected position of the 5’ edge of an exon-junction complex (A, C) or distance between the 5’ edge of a ribosome to the first nucleotide, of a stop codon, in its aminoacyl site (B, D).

We then tested whether changes in EJC and CTRD footprints were associated with the deNADding activity or serine-rich NTD of Arabidopsis DXO1. To do this, we utilized transgenic lines expressing GFP-fused DXO1 variants under the control of the cauliflower mosaic virus-derived 35S promoter (*pro35S*::*DXO1*/*dxo1-2*) (1). This included DXO1(WT), DXO1(E394A/D396A), and DXO1(ΔN194), which represent the Col-0 DXO1 amino acid sequence, DXO1 with amino acid substitutions at the catalytic sites required for deNADding, and deletion of the serine-rich NTD, respectively. We first ensured the integrity of these lines by confirming that they resembled their reported phenotypes (1). Indeed, DXO1(WT) and DXO1(E394A/D396A) resembled WT plants whereas DXO1(ΔN194) more closely resembled the *dxo1* null mutant (**Supplementary Figure 2A**). We then carried out GMUCT on 12-day-old seedlings for each line. We first compared WT with DXO1(WT) and found that over-expression of variant DXO1 in and of itself impacted EJC and CTRD footprints (**Supplementary Figure 2B-C**). This may reflect altered activity or regulation of ectopically expressed DXO1 with a C-terminally fused GFP. Nonetheless, we reasoned that biologically meaningful comparisons could be made between DXO1(E394A/D396A), DXO1(ΔN194), and DXO1(WT). Indeed, DXO1(WT) and DXO1(E394A/D396A) showed highly comparable footprints. On the other hand, DXO1(ΔN194), recapitulated the elevated EJC footprints and loss of CTRD as observed between WT and *dxo1* seedlings (**Figure 2C-D**). Although, the difference in CTRD between transgenic lines is relatively attenuated. Nonetheless, these results suggest that the serine-rich NTD of Arabidopsis DXO1 conveys a function to this protein in mRNA surveillance.

We then explored transcript-specific EJC-binding and CRTD-regulation by calculating a terminal stalling index (TSI) associated with EJC (TSI_EJC_) and CTRD (TSI_CTRD_) footprints for all mRNAs detected in WT and *dxo1* degradomes. We considered an mRNA to be EJC-bound or CTRD-regulated if TSI_EJC_ > 2 or TSI_CTRD_ > 2, respectively. This approach identified 489 and 1,746 EJC-bound mRNAs in WT and *dxo1* plants, respectively (**Figure 3A**, **Supplementary Table 3**). Of these, 1,423 were detected specifically in *dxo1*, which we categorized as DXO1-associated EJC-bound mRNAs (I, **Supplementary Table 4**). We also identified 993 and 389 mRNAs as CTRD-regulated in WT and *dxo1* plants, respectively (**Figure 3B**, **Supplementary Table 5**). Of these, 720 were detected only in WT, which we categorized as DXO1-associated CTRD-regulated mRNAs (II, **Supplementary Table 6**). We compared the magnitude of change in TSI_EJC_ and TSI_CTRD_ between WT and *dxo1* plants. In the case of EJC stalling, *dxo1* showed a statistically significant 2.5× increase in median TSI_EJC_ (**Figure 3C**). Conversely, with respect to CTRD, *dxo1* plants exhibited a statistically significant 2.2× decrease in median TSI_CTRD_ (**Figure 3D**). It is worth noting that, in both cases, there was substantial transcript-specific variation. To test for an association between DXO1-associated EJC-binding, CTRD regulation, and deNADding, we overlapped those mRNAs identified in this study with 1,506 nuclear-encoded NAD^+^-capped mRNAs (out of 1,548 NAD^+^-capped RNAs) previously detected in *dxo1* seedlings (9). Strikingly, each phenomenon occurred on distinct populations of transcripts (**Figure 3E**). We explored which biological processes were associated with each category of transcript using GO enrichment analysis (**Figure 3F**). While category I mRNAs (EJC-bound mRNAs) were related to metabolic processes and gene expression, including translation and RNA processing, category II (CTRD mRNAs) contained terms related to development and signaling. Meanwhile, deNADding targets of DXO1 were enriched for proteins-associated with translation, including a suite of ribosomal proteins, aligning with prior observations (9). Taken together, loss of DXO1 abrogates translation-dependent mRNA surveillance in a deNADding-independent manner.

**Figure 3.**
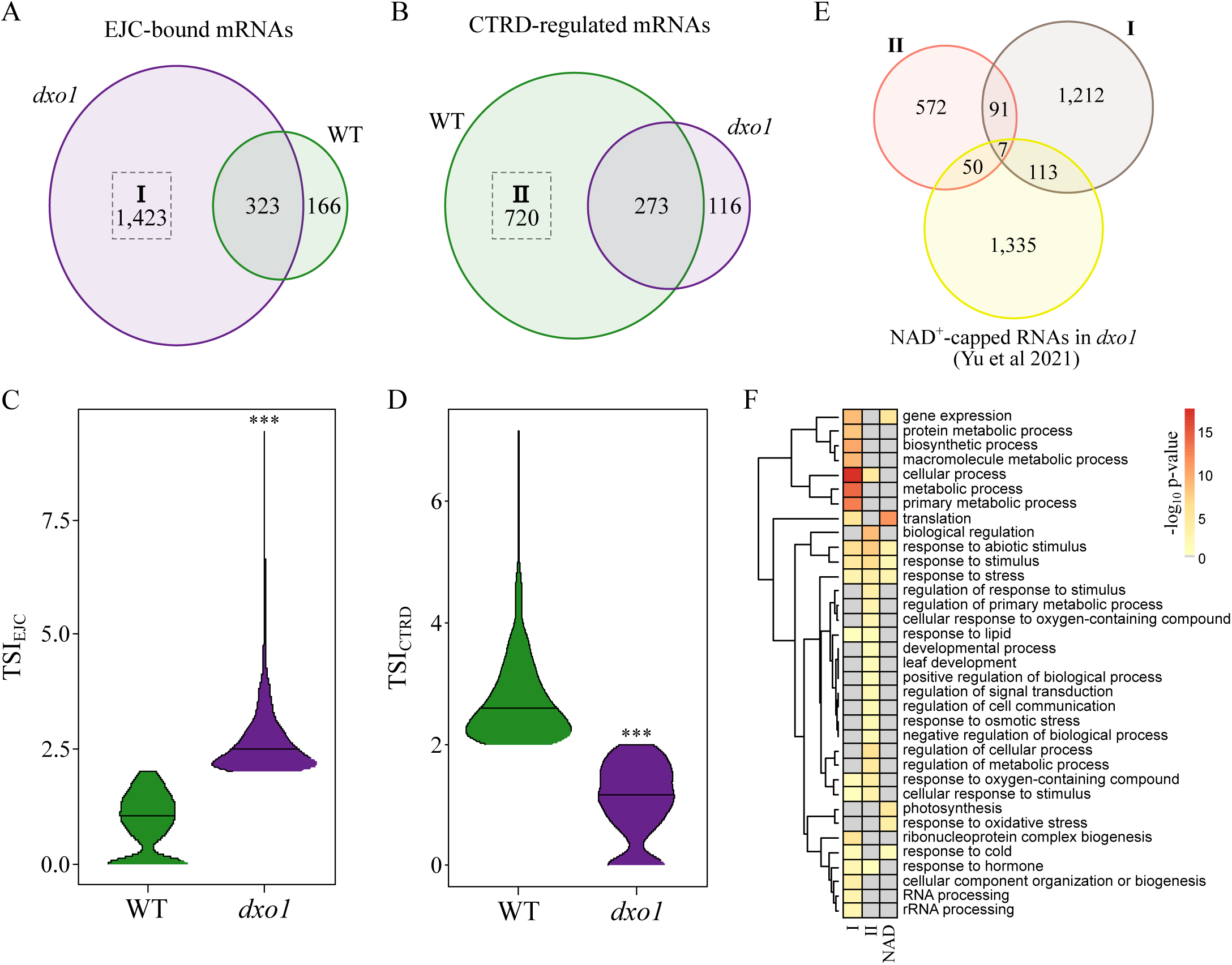
Distinct populations of transcripts show altered EJC footprints, CTRD, and NAD^+^-capping in *dxo1* seedlings. (A) Number of mRNAs harboring stalled EJCs in WT and *dxo1* as determined by TSI_EJC_ > 2. (B) Number of mRNAs actively undergoing CTRD in WT and *dxo1* as determined by TSI_CTRD_ > 2. (C-D) Violin plots of (C) TSI_EJC_ and (D) TSI_CTRD_ in WT and *dxo1* for mRNAs with stalled EJCs (I) and impaired CTRD (II) in *dxo1* plants, respectively. Horizontal line denotes the median. Statistical signifiance was determined using Dunn’s test with Bonferroni-adjusted p-values: *, p < 0.1; **, p < 0.01; ***, p < 0.001. (E) Overlap of mRNAs with stalled EJCs (I), impaired CTRD (II), or that are NAD^+^-capped in *dxo1* seedlings (Yu et al. 2021). (F) Heatmaps with one-dimensional hierarchical clustering of significantly enriched GO terms in mRNAs with stalled EJCs (I) and impaired CTRD (II). Cell colour denotes statistical significance (-log_10_ p-value; grey represents non-significant terms).

### DXO1 impacts pre-rRNA processing and ribosome loading

That many deNADding targets of DXO1 encode ribosomal proteins potentiates defective ribosome biogenesis. Consistent with this are the reported pre-rRNA processing defects in *dxo1* plants, evident by the accumulation of ITS2-first pathway intermediates (11). These observations led us to investigate pre-rRNA processing by mapping our GMUCT reads against a 45S rDNA reference (28). A global view of pre-rRNA processing highlighted elevated cleavage at B_1_ in *dxo1* and DXO1(ΔN194) compared to WT and DXO1(WT), respectively (**Supplementary Figure 3**). This suggests increased processing via the major ITS1-first pathway, involving endonucleolytic cleavage at A_3_ followed by exonucleolytic trimming by XRN2/3 (29). Closer inspection indicated increased cleavage at A_3_ in *dxo1* and DXO1(ΔN194) compared to their respective controls (**Figure 4A**). Although a smaller difference was observed at A_3_ compared to B_1_, likely reflecting rapid processing by XRN2/3. Beyond this, decreased 5′-P ends were observed upstream at the P, P_1_, and P′ sites, as well as within ITS2. These observations suggest that pre-rRNA processing is altered by loss of DXO1 or its serine-rich NTD. In particular, we observed reduced 5′-ETS-first processing, reduced ITS2-first processing, and increased ITS1-first processing. Interestingly, DXO1(E394A/D396A) demonstrated increased processing at P_2_ and within ITS2, while having reduced processing at the A_3_ site. This suggests that DXO1 deNADding impacts pre-18S rRNA processing whereas the serine-rich NTD affects pre-27S rRNA processing.

**Figure 4.**
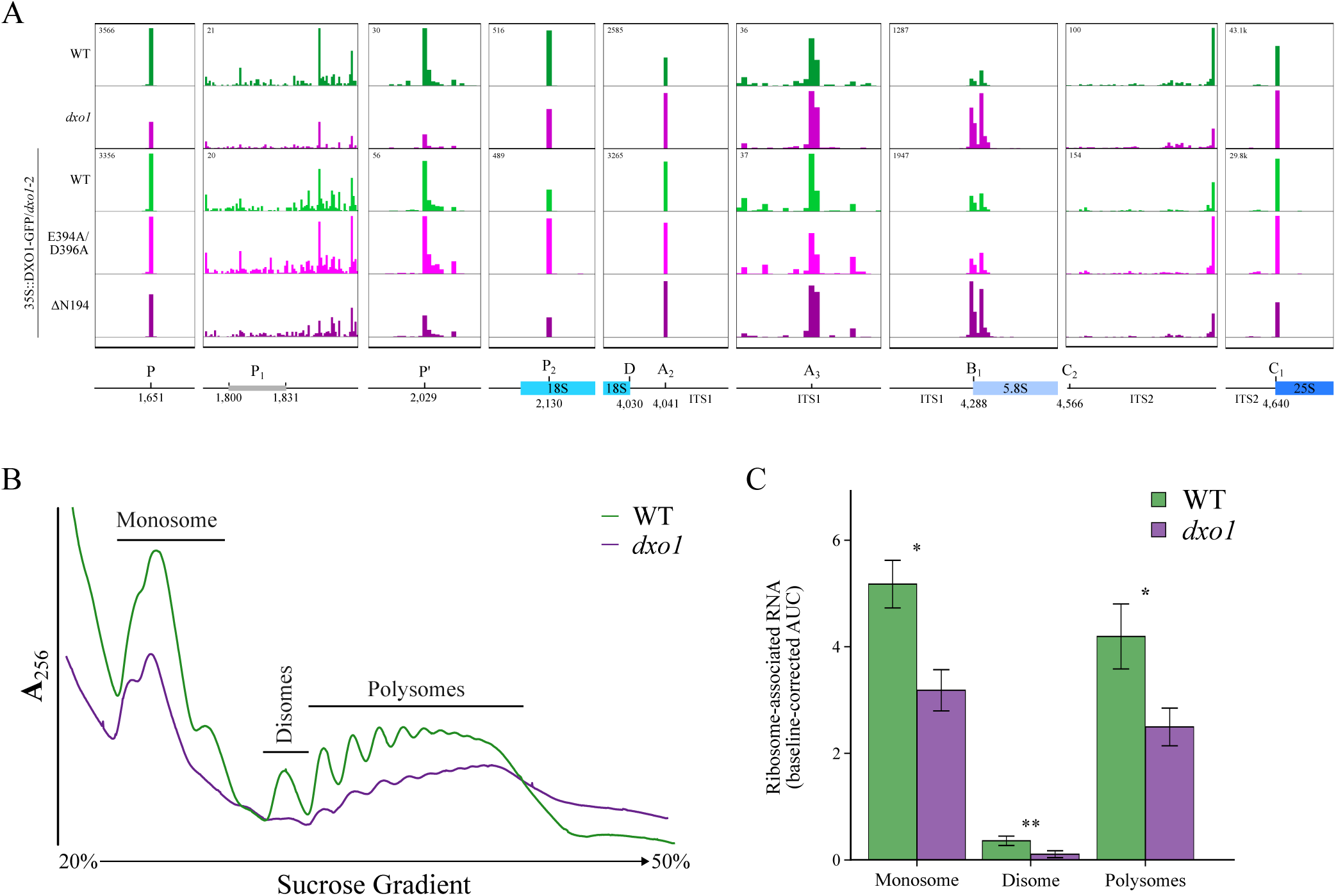
DXO1 influences pre-rRNA processing and ribosome loading in Arabidopsis seedlings. (A) Genome browser views of 5’-P ends at select pre-rRNA cleavage sites in WT, *dxo1*, DXO1(WT), DXO1(E394A/D396A), and DXO1(Δ194) degradomes. Numbers reflect the position of the indicated cleavage sites in the rDNA reference sequence (Sims et al 2019). Cleavage site positions are based on Sáez-Vásquez and Delseny (2019). B) Representative polysome profiles based on absorbance at a wavelength of 256 nm (**A**_256_) of ribosome extracts from WT and *dxo1* seedlings separated by sucrose gradient fractionation. The regions of the gradient pooled into monosome, disome, and polysome are indicated. C) Quantification of ribosome-associated RNA content in each genotype, based on the baseline-corrected area under the curve (AUC), for the specified fractions identified in the polysome profiling. Significance was determined using Tukey’s test of multiple comparisons: * p < 0.05, ** p < 0.01.

Next, we considered the possibility that pre-rRNAs can be NAD^+^-capped and targeted by DXO1. To do this, we re-analyzed total RNA-sequencing (RNA-seq) and NAD^+^-capped RNA-sequencing (NAD-seq) performed on WT and *dxo1* seedlings (GSE142388, **Supplementary Table 2**) (9). Importantly, both library preparations facilitated the capture of non-polyadenylated RNAs, which were mapped to the 45S rDNA assembly to calculate the proportion of NAD^+^-capped pre-rRNAs. In general, we observed a low proportion of NAD^+^-capped pre-rRNAs except for 18S, which was enriched relative to total RNA (**Table 2**). Nevertheless, they could be detected across the 45S rDNA assembly and we observed increases for ITS1, 5.8S, and ITS2, and a decrease for 25S. Further investigation revealed that these changes were driven by changes in NAD-seq samples (**Table 3**). The largest change in proportion NAD^+^-capped pre-rRNAs was derived from ITS1, which also had the most pronounced change in pre-rRNA processing (ITS1-first pathway, **Supplementary Figure 3A**). However, this altered processing appears to be deNADding-indendent. Nevertheless, this analysis suggests that pre-rRNAs can be NAD^+^-capped (at low levels) and targeted by DXO1.

**Table 2.** Proportion NAD^+^-capped pre-rRNAs in WT and *dxo1* seedlings.

| Pre-rRNA | Proportion NAD<sup>+</sup>-RNA<br>(NAD-seq<sub>RPM</sub>/RNA-seq<sub>RPM</sub>) |  |  |  | Fold-change<br>(*dxo1*/WT) |
| --- | --- | --- | --- | --- | --- |
|  | WT |  | *dxo1* |  |  |
|  | Mean | SE | Mean | SE |  |
| 18S | 1.93 | 0.20 | 1.93 | 0.12 | 1.00 |
| ITS1 | 0.11 | 0.003 | 0.24 | 0.07 | 2.07 |
| 5.8S | 0.23 | 0.03 | 0.34 | 0.06 | 1.46 |
| ITS2 | 0.61 | 0.10 | 0.69 | 0.09 | 1.13 |
| 25S | 1.20 | 0.006 | 1.03 | 0.09 | 0.86 |

**Table 3.** Relative abundance of total and NAD^+^-capped pre-rRNAs between WT and *dxo1* plants.

| Pre-rRNA | RNA-seq |  | NAD-seq |  |
| --- | --- | --- | --- | --- |
|  | Fold-change<br>(*dxo1*/WT) | FDR-adjusted<br>p-value | Fold-change<br>(*dxo1*/WT) | FDR-adjusted<br>p-value |
| 18S | 0.97 | 0.960 | 0.98 | 0.663 |
| ITS1 | 0.91 | 0.960 | 1.66 | 0.043 |
| 5.8S | 1.01 | 0.960 | 1.47 | 0.003 |
| ITS2 | 1.65 | 0.960 | 1.92 | 0.046 |
| 25S | 0.98 | 0.960 | 0.85 | 0.015 |

The observed changes in pre-rRNA processing and NAD^+^-capped pre-rRNAs could result in, or reflect, changes in ribosome function. We performed polysome profiling to assess changes in mRNA-ribosome associations. This was performed in biological triplicate, resulting in reproducible profiles for each genotype (**Supplementary Figure 3B**). Analysis of representative profiles revealed a striking depletion for all populations of ribosomes in *dxo1* compared to WT seedlings (**Figure 4B**). To quantify the difference, we determined the baseline-corrected area under the curve for regions representing a single RNA-bound ribosome (monosomes), two ribosomes (disomes), and three or more ribosomes (polysomes) for each genotype (**Figure 4C**). From this, we estimate that *dxo1* seedlings had 61.5%, 29.6%, and 59.5% of the monosome-, disome-, and polysome-associated RNA content as compared to WT, respectively, indicating that DXO1 is critical for ribosome loading. Taken together, these results suggest that DXO1 is important for rRNA processing and ribosome loading.

## Discussion

In this study, we investigate the evolution of Arabidopsis DXO1 and its impacts on mRNA surveillance and translation. We show that Arabidopsis DXO1 is highly diverged from other homologs, including those in non-flowering plants, and attribute this to the presence of a serine-rich NTD that is likely limited to Angiosperms. Degradome analysis revealed that loss of this domain results in abrogated EJC clearance and CTRD for distinct populations of transcripts but does not correlate with transcripts that are deNADded by DXO1. Lastly, *dxo1* seedlings exhibit abrogated pre-rRNA processing and ribosome loading. Collectively, we propose that Arabidopsis DXO1 is an evolutionarily divergent homolog that impacts ribosome loading, which is required for translation-dependent mRNA surveillance.

Our evolutionary analysis suggests that angiosperms harbor a unique DXO1 homolog that has diverged multiple times. Lack of a prokaryotic root makes it difficult to discern the exact ancestry. However, we hypothesize that there was a shared DXO lineage followed by the divergence of plant DXO1, then mammalian DXO from fungal Rai1 (**Figure 1E**). This is consistent with the biochemical uniqueness of Arabidopsis DXO1, which is primarily a deNADding enzyme with limited activity against other 5′-cap structures, including 5′-m^7^G, 5′-PO_4_, and 5′-OH (1, 2). The loss of 5′-decapping and exoribonuclease activity, a hallmark of mammalian and fungal DXO homologs, appears to result from a plant-specific asparagine residue (Asn298) within the expected RNA binding site (1). In addition, we observe another divergence between angiosperms and non-flowering plants. Our analysis suggests that the serine-rich NTD of Arabidopsis DXO1 likely arose in angiosperms (**Figure 1D**). This region is required for activating RNMT1 to catalyze 5′-cap methylation (12). The molecular importance of this interaction remains unclear. One hypothesis is that DXO1 recruits RNMT1 to non-canonically capped RNAs, including NAD^+^-capped RNA, to facilitate 5′-m^7^G recapping. This may represent an angiosperm-specific form of mRNA surveillance leading to recapping, whereas fungal Rai1 and mammalian DXO degrade non-canonically capped RNA (3, 7). This raises questions surrounding DXO1 function in non-flowering plants that do not harbor this region (**Figure 1C**). Degradome profiling in these species lacking an Arabidopsis-like DXO1 homolog would help establish whether the serine-rich NTD confers such a specialized function.

Both EJC and CTRD footprints reflect the activity of translation-dependent mRNA surveillance, that is, they are indicative of ribosome activity (18–20). We find that these footprints are altered in plants lacking DXO1, indicating defective mRNA surveillance (**Figure 2**). For instance, elevated EJC footprints likely reflect increased EJC-bound mRNAs unable to undergo the pioneer round of translation. Meanwhile, CTRD may be suppressed since it occurs in the wake of an elongating ribosome, which requires ribosome loading (**Figure 4B**). That these footprints are altered in DXO1(ΔN194) plants (**Figures 2C-D**) implicates RNMT1, or other interactors, to this process. A recent study posits that deNADding by DXO1 contributes to CTRD of NAD^+^-capped transcripts (30). However, our findings question this notion. Firstly, deNADding targets of DXO1 (9), which were not considered previously and do not reflect the entire NAD^+^-capped RNA pool, did not show altered EJC or CTRD footprints in *dxo1* degradomes (**Figure 3E**). Secondly, we highlight that DXO1(E394A/D396A) and DXO1(ΔN194) must be compared to the transgenic WT control [DXO1(WT)], rather than to wild-type Col-0 as was done previously (30), since the control line already demonstrates suppressed CTRD (**Supplementary Figure 2**). By incorporating DXO1(WT), we reveal that the serine-rich NTD is more relevant for mRNA surveillance (including CTRD) than deNADding (**Figures 2C-D**). We propose that the loss of mRNA surveillance-triggered decay in *dxo1* mutant plants is the consequence of deficient ribosome loading, the precursor event (**Figure 4B**). What remains unclear is the cause of defective ribosome loading. The presence of the canonical 5′-m^7^G-cap is critical for recognition by m^7^G cap-binding proteins, including the nuclear cap-binding complex and eIF4E that facilitate ribosome loading for the pioneer round of translation and steady-state translation, respectively (31, 32). Therefore, a possible culprit is RNMT1 inaction since it is activated by DXO1 for 5′-cap methylation (12). Indeed, co-activation of mammalian RNMT-RAM in activating T cells was found to promote ribosome biogenesis (33). Together, this warrants investigation of RNMT1 involvement in mRNA surveillance and ribosome loading. Furthermore, the nature of the substrates on which DXO1-RNMT1 operate, likely extending beyond NAD^+^-capped RNAs, also remains to be determined.

Plants lacking DXO1 demonstrated changes in pre-rRNA processing (**Figure 4A**, **Supplementary Figure 3**) and NAD^+^-capped pre-rRNA levels (**Tables 2**–**3**). This is similar to the Rai1 paralog in *Saccharomyces cerevisiae*, Dxo1, which acts as a distributive exonuclease to facilitate 5′-end processing of 25S rRNA (8). However, it is unclear whether Arabidopsis DXO1 is directly involved in pre-rRNA processing given its limited exonucleolytic activity (1). Instead, altered processing can be attributed to the serine-rich NTD as shown here and previously (11). Since rRNAs lack the m^7^G-cap, it is likely that RNMT1 influences pre-rRNA processing indirectly. For example, in human cell lines, RNMT-RAM influences 45S rRNA synthesis by impacting RNA polymerase I activity (34). Whether Arabidopsis DXO1-RNMT1 shares this function remains to be investigated. Additionally, in yeast, secondary structure at the 5′-terminus of 5S rRNA is critical for 5S rRNA-protein complex formation (35). Therefore, it may also be worthwhile investigating whether Arabidopsis DXO1 has additional interacting partners that could impact rRNA-protein associations. Nevertheless, altered pre-rRNA processing co-occurred with reduced ribosome loading (**Figure 4B**). That the loss of DXO1 results in phenotypes (9) resembling plants lacking ribosome biogenesis factors (36) or ribosomal proteins (37) suggests that the root cause of abnormalities in *dxo1* null mutants are due to deficient ribosome assembly and/or function. In this case, the observed changes in pre-rRNA processing likely reflects feedback regulation. For example, pre-60S maturation is a prerequisite for pre-40S maturation, whereby impaired pre-60S maturation results in the accumulation of 18S rRNA precursors (38). However, whether increased ITS1-first processing in *dxo1* seedlings reflects a compensatory adjustment to maintain ribosomal subunit stoichiometry remains to be elucidated.

In conclusion, we demonstrate that Arabidopsis DXO1 is an evolutionarily diverged homolog that contributes to ribosome loading and mRNA surveillance. The extent to which homologs in mammals, fungi, or non-flowering plants retain this function remains unknown. The deNADding activity of Arabidopsis DXO1 is inhibited by 3′-phosphoadenosine 5′-phosphate (PAP) (1), a chloroplast-derived retrograde signal that contributes to oxidative stress responses (39, 40). Based on our observations, this provides a mechanism for retrograde signalling to directly regulate ribosome loading, although it is unclear whether PAP inhibits the deNADding-independent function of DXO1. Consistent with this hypothesis is that *dxo1* plants are stress sensitive and show differential temperature-responsive regulation of pre-rRNA processing (11). This raises new questions regarding the relative contributions of PAP-dependent inhibition of plant XRNs (41) and DXO1 (1) towards plant acclimation and adaptation that should be addressed in the future.

## Materials and Methods

### Evolutionary analysis

The amino acid sequence representing the primary isoform of Arabidopsis DXO1 (AT4G17620.1) was used to identify its cognate Pfam clan (CLO236) using HmmScan (42). Sequences within Pfam clan CLO236 were used to generate a sequence similarity network using EFI-EST to generate nodes with ≥40% sequence similarity (43). The node containing AT4G17620.1 was used to attempt to identify a possible prokaryotic common ancestor; which was not found. Therefore, the Pfam family PF08652 was used for further analyses. These sequences underwent manual curation to incorporate taxonomic information (BioPython Entrez) and to remove duplicate sequences or those with ambiguous amino acids. Sequences of length >2 standard deviations of the mean were removed. Sequences of Kingdoms with fewer than 30 orthologs were also removed. Evolutionary analysis was then performed on this filtered, high-confidence dataset (**Supplementary Figure 1A**, **Supplementary Table 1**).

The presence of a serine-rich NTD was systematically determined using the frequency of serine residues within the first 18% of amino acids for each homolog, based on manual curation of sequence alignments performed with T-COFFEE using default settings (44). A threshold frequency of 0.23 was determined, based on the trough in its bimodal distribution, above which we considered to reflect the presence of this domain (**Supplementary Figure 1C**). Sequences with fewer than 10 serine residues were excluded to avoid artifacts caused by short sequences. Cladograms were based on hierarchical relationships between taxa defined in Pfam, with some modification: Zygnematophyceae was placed as a sister group to Embryophyta, mosses and liverworts were grouped together, and Amborellale was moved to be a sister to all other angiosperms, followed by Nymphaeales. Pseudotime analysis was performed in Python 3.8 using the Evolocity pseudotime method with the esm1b model (45), Uniform Manifold Approximate Projection (UMAP) dimensionality reduction at a scale of 1.0, and nearest-neighbor clustering at 25 sequences (25).

### Plant material

All *Arabidopsis thaliana* (L.) (Arabidopsis) germplasm studied were in the Columbia (Col-0) background. Arabidopsis lines include wild-type Col-0 (WT) and *dxo1* (*dxo1*-2, SALK_032903), which were obtained from the Arabidopsis Biological Resource Centre (Ohio State University, Columbus, OH, USA). Homozygous *dxo1* mutants were identified based on PCR amplification of SALK_032903 (and absence of amplification with gene-specific primers flanking the predicted insertion site). The DXO1(WT), DXO1(E394A/D396A), and DXO1(ΔN194) transgenic lines (1) were kindly provided by the Kufel lab (University of Warsaw, Warsaw, Poland). Individuals harboring a *pro35S::DXO1-GFP* transgene were identified by PCR amplification. Primers used for PCR are listed in **Supplementary Table 7**. Progeny collected by single-seed descent from homozygous parents were used for experiments after ensuring that there was no segregation of the expected phenotypes (1, 9).

### Plant growth

Arabidopsis seedlings were grown on half-strength Murashige and Skoog medium (Phytotech Labs, Lenexa, KS, USA) at 22 °C under a 16-h-light (100-150 µmol m^-2^ s^-1^)/8-h-dark photoperiod. For soil-based growth, seeds were sown onto moist soil (treated overnight with diatomaceous earth), supplemented with 10 g/L Jack’s Professional 15-16-17 Peat-Lite water-soluble fertilizer (JR Peters, PA, USA) and NemAttack (Arbico Organics, AZ, USA), then stratified in the dark for 48 h at 4 °C. Plants were grown at 22 °C under a 16-h-light (100-150 µmol m^-2^ s^-1^)/8-h-dark photoperiod.

### GMUCT

The degradome was assessed using GMUCT, which was performed on 12-day-old plate-grown seedlings. Tissue was snap-frozen in liquid N_2_ and homogenized with an autoclaved and chilled mortar and pestle. Cells were lysed using Qiazol lysis reagent (Qiagen, Valencia, CA, USA) and the lysate was homogenized using a Qiashredder (Qiagen). Total RNA was then isolated using Qiagen miRNEasy mini columns (Qiagen) as per the manufacturer’s instructions. The RNA was treated with RNase-free DNase (Qiagen) at room temperature for 30 minutes followed by purification via ethanol precipitation. The purified RNA was used to prepare GMUCT libraries using an established protocol (17). Libraries underwent single-end sequencing (50 cycles) on an Illumina HiSeq2000.

Raw GMUCT reads were trimmed using TrimGalore (Babraham Bioinformatics) to remove adapter sequences, low-quality base calls (PHRED < 20), and sequences shorter than 10 nucleotides (--length 10). Trimmed reads were aligned to the Arabidopsis TAIR10 reference using STAR (46) with flags --outFilterMismatchNmax 0 --outFilterScoreMinOverLread 0.75 --outFilterMatchNminOverLread 0.75 --outFilterMultimapNmax 1. For assessing pre-rRNA processing, reads were mapped against an rDNA reference sequence (variant 1) (28). This was annotated based on homology to characterized rRNA sequences (47, 48) and defined cleavage sites (29), with some adjustments made based on observed 5′-P ends near the following endonucleolytic cleavage sites: P, P′, P_2_, and A_3_.

EJC and CTRD footprints were assessed by summarizing monophosphorylated RNA (5′-P) end counts: (1) upstream of exon-exon junctions (for exons of length ≥ 50 nucleotides) for EJC footprints, and (2) upstream of stop codons for CTRD footprints (18, 19). These regions were only considered for nuclear-encoded protein-coding mRNAs. Aligned reads were assigned a genomic location based on the first nucleotide of each sequence, and the read depth at this position (5′-P end counts) was calculated using Bedtools (genomecov -5 -scale) (49). 5′-P end counts were scaled by the total number of mapped reads, as determined by samtools (samtools view -F 260 -c), and expressed as reads per million (RPM). The relative frequency of 5′-P ends was calculated using 5′-P end counts as previously described (18). First, normalized 5′-P end counts (*P_i_*) were calculated by:

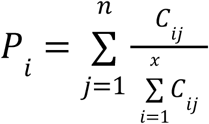

where *C_ij_* is the 5′-P end count at position *i* of region *j*, *x* is the length of the region, and *n* is the number of analyzed regions with scaled 5′-P end count ≥ 1. Then, the relative frequency of 5′-P ends was calculated by dividing *P_i_* at each position by the sum of *Pi* across that region.

To consider whether an mRNA was undergoing co-translational RNA decay, we calculated its Terminal Stalling Index (TSI) (50). TSI_CTRD_ was calculated using 5′-P end counts occurring in a 50 nucleotide region upstream of stop codons:

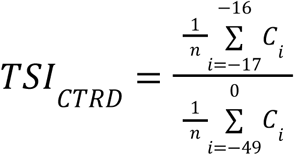

where *C_i_*is the 5′-P end count at position *i* relative to the first nucleotide of the stop codon (position 0) and *n* is the number of positions with at least one raw 5′-P end count. Nucleotides at positions -17 and -16 represent the 5′-edge of a terminating ribosome based on the peak signal observed.

To identify mRNAs harboring stalled EJCs, TSI was calculated based on EJC footprints (TSI_EJC_) using a modified TSI formula:

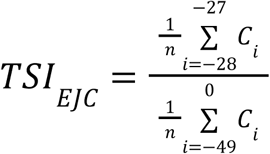

where *C_i_* is the 5′-P end count at position *i* relative to the exon-exon junction (position 0) and *n* is the number of positions with at least one raw 5′-P end count. Nucleotides at position -28 and -27 represent the 5′-edge of the EJC based on the peak signal observed.

Transcripts with mean TSI_CTRD_ or TSI_EJC_ ≥ 2 and raw 5′-P end count ≥ 10, were considered active targets for CTRD or EJC stalling, respectively, for each sample group. Enriched GO terms were determined using the statistical overrepresentation test (Fisher’s exact test with Bonferroni correction) from PANTHER (51). Only GO terms with a fold enrichment > 1, in at least one sample group, were considered and redundancy was removed using rrvgo (52).

### Polysome profiling

Polysome profiling was performed on 14-day old plate-grown seedlings using an adapted protocol (53). Briefly, 200 mg of seedlings were homogenized in 800 µl of extraction buffer (160 mM Tris-Cl [pH 7.6], 80 mM KCl, 5 mM MgCl_2_, 5.36 mM EGTA [pH 8], 0.5% IGEPAL CA-630, 40 U ml^-1^ RNasin Plus RNase inhibitor [Promega], 150 µg ml^-1^ cycloheximide and 150 µg ml^-1^ chloramphenicol) and incubated on ice for 10 min. Samples were repeatedly centrifuged at 16,000 g for 5 min at 4 °C until the supernatant was clear. 700 µl of supernatant was loaded onto a sucrose gradient, consisting of layers of 50% (1.68 ml), 35% (3.32 ml), 20% (3.32 ml) and 20% (1.68 ml), with the two 20% layers added separately. The buffer used in the gradients consisted of 400 mM Tris-Cl (pH 8.4), 200 mM KCl, 100 mM MgCl_2_, 10.12 µg ml^-1^ cycloheximide and 10.12 µg ml^-1^ chloramphenicol. Gradients were centrifuged at 36,000 g with an SW41Ti rotor for 3.5 h at 4 °C. Each gradient was passed through a spectrophotometer from the highest to lowest density and absorbance at 256 nm was recorded, with 1 ml fractions collected using a Gradient Piston Fractionator (BioComp). Sucrose gradient fractions were collected based on the observation of peaks representing 1 ribosome (monosomes), 2 ribosomes (disomes), and 3 or more ribosomes (polysomes, **Figure 4B**). Total RNA was isolated from each fraction using 5:1 (v/v) acid phenol:chloroform. Briefly, an equal volume of the phenol:chloroform mix was added to each fraction, before centrifugation at 16,000 g for 10 min. The aqueous layer was extracted a second time using a one-fifth volume of chloroform, before the RNA was precipitated using an equal volume of isopropanol. The precipitated RNA was washed twice with 70% ethanol and resuspended in nuclease-free water. Five micrograms of purified RNA was treated with TURBO DNase (Thermo Fisher Scientific, Waltham, MA, USA) for 30 min at 37°C. DNA-free RNA was purified using Sera-Mag paramagnetic beads (1.8× bead:sample). RNA was quantified with a Qubit 2.0 Fluorometer (Thermo Fisher Scientific), and RNA quality was assessed via a Nucleic Acid Analyzer LabChip GX Touch (PerkinElmer, Shelton, CT, USA).

## Data availability

Sequencing data generated is available at NCBI Gene Expression Omnibus under accession GSE285834. Published sequencing data was accessed from GSE71913 and GSE142388. Descriptions of sequencing data analyzed are provided in **Supplementary Table 2**. Files containing the 45S rDNA reference sequence and annotation are available at figshare DOI: 10.6084/m9.figshare.30471215.

## Acknowledgements

We are grateful to Joanna Kufel for kindly providing seeds of the DXO1(WT), DXO1(E394A/D396A), and DXO1(ΔN194) transgenic lines. We also thank Benjamin Wales-McGrath for assistance with screening Arabidopsis lines.

## Author contributions

DRG, XY, and BDG designed the study. XY performed GMUCT. MDM performed evolutionary analysis. MM and PK performed polysome profiling. DRG analyzed the datasets and wrote the manuscript. All authors reviewed and commented on the manuscript.

---

**Supplementary Figure 1.**
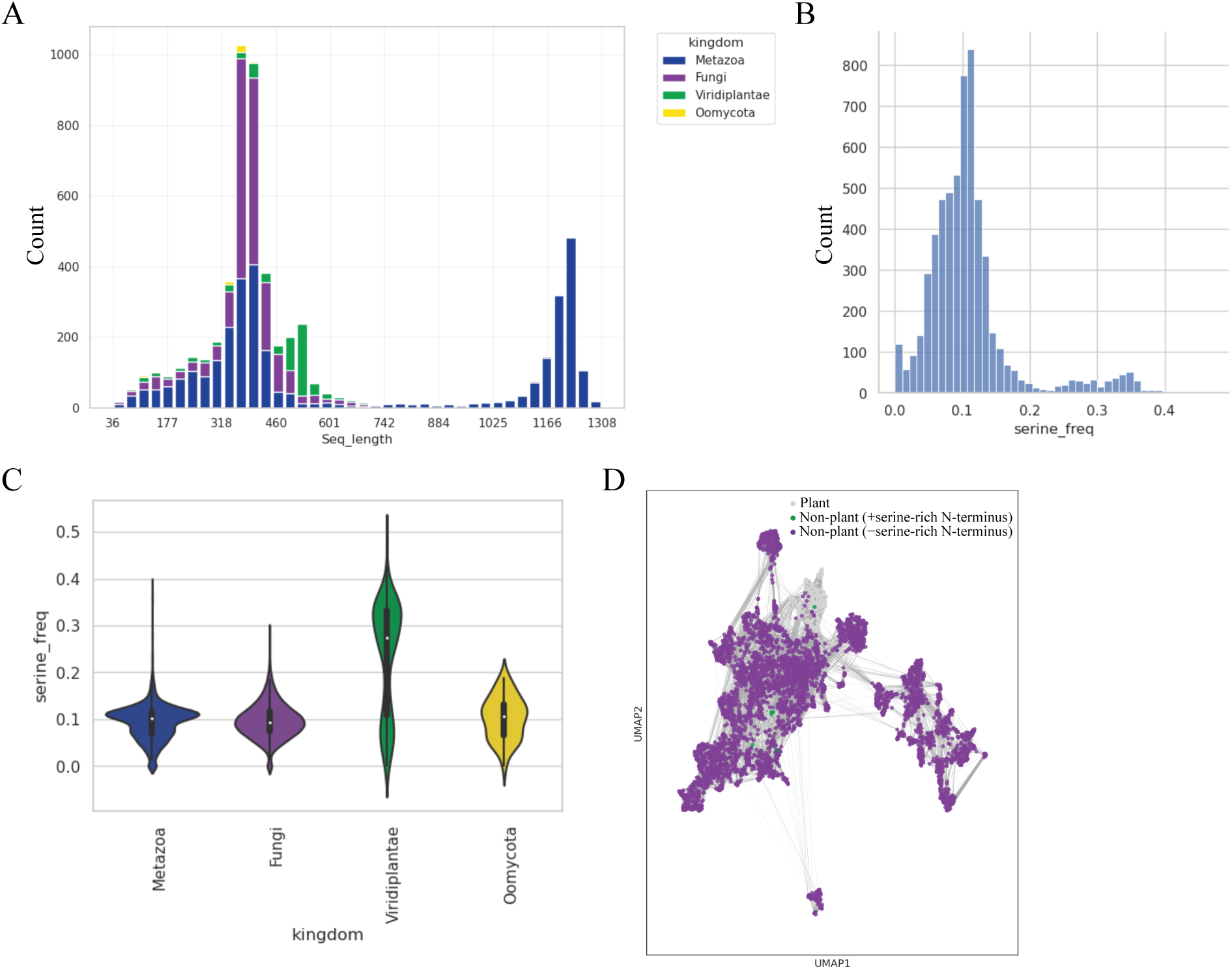
Sequence information for Rai1/Dxo1/DXO homologs. (A) Length distribution of curated and filtered sequences from Pfam family PF08652 comprising the dataset used for evolutionary analysis. Bar colour denotes kingdom. (B) Serine frequency distribution within the first 18% of sequences in our dataset. (C) Violin plot of serine frequency distribution within the first 18% of sequences in our dataset grouped by kingdom. (D) KNN network from Figure 1 B recolored to highlight non-plant sequences with (green) and without (purple) a serine-rich N-terminus. Plant sequences are shown in grey.

**Supplementary Figure 2.**
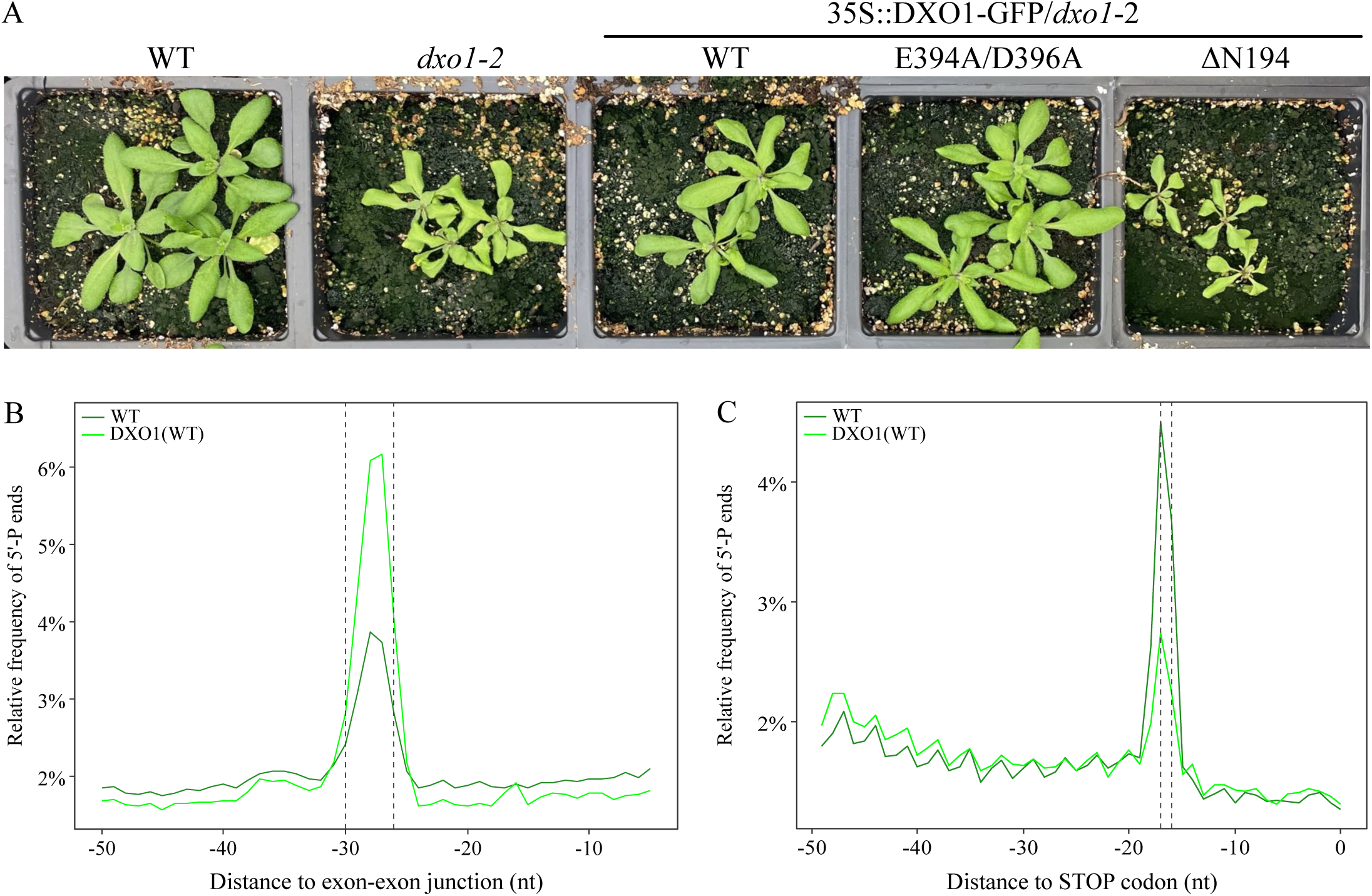
Overexpression of DXO1 fused to GFP impacts RNA turnover in Arabidopsis. (A) Representative phenotypes of three-week old soil-grown WT, *dxo1*, and transgenic DXO1 (35S::DXO1/*dxo1*-2, Kwasnik et al 2019 Nucleic Acids Research) plants. (B) Relative frequency of 5’-P ends upstream of exon-exon junctions in WT and DXO1(WT). Dotted lines represent the expected position of the 5’ edge of an EJC. (C) Relative frequency of 5’-P ends upstream of stop codons in WT and DXO1(WT). Dotted lines represent the distance between the 5’ edge of a ribosome to the first nucleotide, of a stop codon, in its aminoacyl site.

**Supplementary Figure 3.**
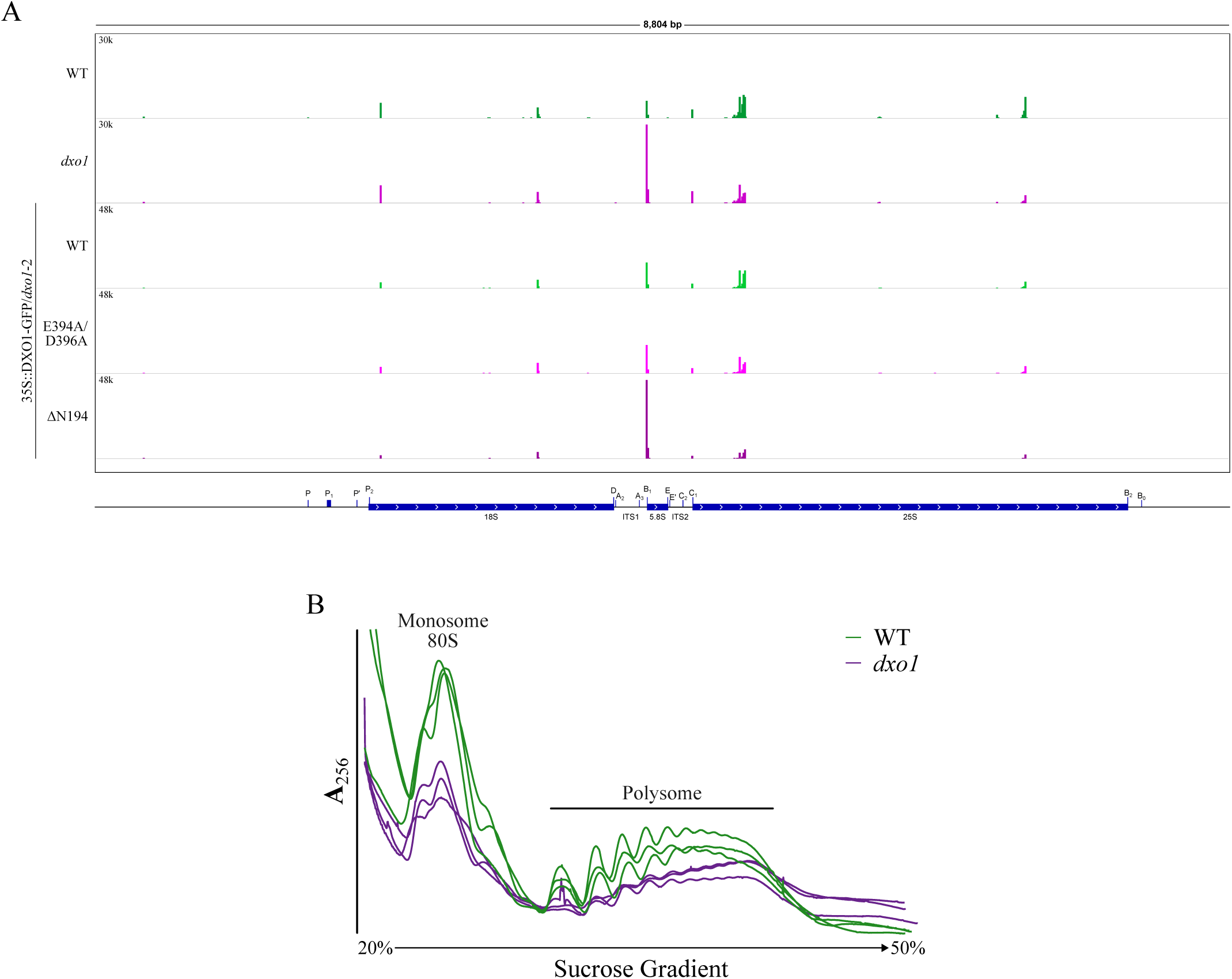
Assessing pre-rRNA cleavage and polysome profiles in seedlings lacking DXO1. (A) Genome browser view of 5’-P ends across 45S pre-rRNA in WT, *dxo1*, DXO1(WT), DXO1(E394A/ D396A), and DXO1(Δ194) degradomes. (B) Three biological replicates of polysome profiles based on absorbance at a wavelength of 256 nm (**A**_256_) of ribosome extracts from WT and *dxo1* seedlings separated by sucrose gradient fractionation.

